# Genomewide analysis of biomass responses to water withholding in young plants of maize inbred lines with expired plant variety protection certificate

**DOI:** 10.1101/704668

**Authors:** Maja Mazur, Andrija Brkić, Domagoj Šimić, Josip Brkić, Antun Jambrović, Zvonimir Zdunić, Vlatko Galic

## Abstract

Patterns of seasonal variations in rainfall are changing which affects rain-fed agricultural areas. Water deficit during the early vegetative growth poses threat as it causes variability in plant development, makes plant susceptible to other stresses and deteriorates the stands. Plant’s responses to water deficit are reflected in biomass traits which represent the morpho-physiological adjustments of plant to new conditions. The aims of this study were to assess the biomass responses of maize inbred lines with expired plant variety protection certificate that are freely distributed worldwide using genomewide analysis approach. The collection of 109 maize inbred lines, genotyped using genotyping-by-sequencing, was planted in controlled conditions (16/8 day/night, 25°C, 50% RH, 200 μMol/m^2^/s) in trays filled with soil in three replicates. Plants in control (C) were watered every two days with 8 ml_H2O_, while watering was stopped for 10 days in water withholding (WW) treatment. Fourteen days old plants were harvested and fresh weight (FW), dry weight (DW) and dry matter content (DMC, % of FW) were measured. Different responses to WW were detected in two genetic subgroups: Stiff Stalk and Non-Stiff Stalk. Totally 29 QTLs were detected, and it was shown that genetic regulation of DMC is different than regulation of FW and DW. This was further supported with correlations of rrBLUP marker effects among the traits. It was concluded that measurements of biomass traits in this manner are fast and reliable indicator of plant’s response to water deficit and can be used for effective screening of breeding progenies.

## 1. Introduction

Due to climate change with unpredictable nature of drought, breeding crops tolerant to abiotic stress has become crucial for the crop production sustainability [1–3]. Water deficit comes as the main abiotic stress in low input rain-fed systems that account for more than 70% of the maize production in the developing world [4]. The global trends in temperature and precipitation are changing resulting in extreme weather events which may occur at any time throughout the year. Changes in seasonal variations of temperatures inevitably lead to changes in seasonal variations of precipitation, due to the very complex pathways of formations of atmospheric states [5]. Monitoring of satellite observations showed patterns in trends of worldwide changes in precipitation leading to drying of many rain-fed agricultural areas in large parts of Europe and Asia [6]. In these changes, it seems that no climatological scenario will stay unaffected [7]. For example, in Southeastern Europe, maize sowing is mostly limited to the April, due to high frost incidence in March, and inability of plants from all maturity groups to finish the vegetative cycle in terms of growth and dry-down if sowing occurs after first few days of May. In this regard, the changes in precipitation during spring are vastly affecting the maize production in all parts of the world affected by climate change, resulting in unequal development of plants or death, consequently leading to higher disease susceptibility, depletion of stands and many other unfavorable conditions [8]. For example, according to the data from Croatian Meteorological and Hydrological Service for city of Osijek located in the region considered as European Corn Belt, during the last four growing seasons (2015 – 2018), in all years except 2017 there was at least ten-day streak with hydrological drought in April. Although there was no hydrological drought in 2017, there was a 12-day period with no rain. Furthermore, in two consecutive years (2015, 2016), precipitation was for ten or more days below the 10^th^ percentile of 50-year reference period (S2 Figure).

Pannonian basin and Danube regions located in Central and Southeastern Europe, where most of the European maize is grown, can be compared to the US Corn Belt [9]. The genetic material developed in Corn Belt can be grown in this region without need for adaptation, and the hybrid breeding is based on combining inbred lines from two major heterotic groups, Stiff Stalk and Non-Stiff Stalk [10]. In the 1980s vast majority of all commercial maize hybrids in the US were developed from six public inbred lines or their descendants [11]. By the mid-2000s only seven inbred lines (two of them are public) were pointed out as a foundation of the whole North American maize germplasm pool [12]. Several inbred lines were detected as the major progenitors of commercial proprietary US germplasm – DK3IIH6, B73, and PH207, respectively [13]. After genotyping a huge collection of inbred lines, from USDA-ARS North Central Regional Plant Introduction Station (NCRPIS, Ames, Iowa), and other seedbanks around the world, Romay et al. [14] provided valuable genotypic information for researchers and maize breeders. Majority of the elite inbred lines developed from the 1980s up to the present were commercial inbred lines under Plant Variety Protection (PVP) Act, usually with expiration period of 18 years. Ex-PVP germplasm is one of the key germplasm resources in maize breeding today.

Since tolerance to water deficit is a quantitative trait, traditional breeding strategies for superior genotypes selection relying on phenotype have proven to be of limited success due to low heritability, interaction genotype-environment and polygenic effects [15]. For example, in the US Corn Belt, despite the cultivar improvements and higher grain yield performance, sensitivity to water deficit remained the same or even increased over the last 18 years [16]. Combined efforts across scientific disciplines (genetics, physiology, plant breeding) are needed in order to improve maize plant’s responses to water deficit. In recent years, genomewide association studies (GWAS) has developed into valuable approach for identifying gene function [17]. The GWAS approach overcomes several limitations of biparental quantitative trait loci (QTL) mapping. QTL mapping in biparental populations is limited in the amount of the gentic variation and the number of recombination events resulting in lower mapping resolution. GWAS provides higher resolution of QTL location and assays a wide range of natural variation [18]. Using available germplasm resources along with the growing array of new methods and techniques in QTL mapping could provide efficient economic and ecological solutions in both maize breeding and germplasm conservation. Regarding the phenotype, the traits assessing the plant biomass, and dry matter content appear to reflect the effects of water withholding on plant level [19]. Furthermore, there is a considerable genotypic variability present for biomass traits, and they well represent the effects of water deficit on plant morpho-physiological status [20]. The aims of this study were to assess the potential of biomass traits in screening of young maize plants responses to water withholding, and to investigate the genetic background of these responses.

## 2. Materials and Methods

### 2.1. Plant material and experimental design

The seeds of 154 maize inbred lines with expired PVP certificates were obtained from USA Department of Agriculture NCRPIS (Ames, Iowa). Seeds were planted in growing season 2018 and selfed to obtain sufficient quantity of seeds for experiments. Selfing was successful for totally 109 inbred lines. The list of 109 inbred lines with data from patents and repositories is available in S1 Table.

Experiments were set in growth chamber (25°C, RH = 50%, 16/8 day/night, 200 μmol/m^2^/s) in trays (510 × 350 × 200 mm). Each tray was filled with 20 l of soil (pH (CaCl_3_) = 5.7, N (NH_4_^+^ + NO_3_^-^) = 70 mg/l, P (P_2_O_5_) = 50 mg/l, K (K_2_O) = 90 mg/l, EC = 40 mS/m) and divided to 15 rows with 7 boxes (50 × 35 mm) each. The experiment was set with single water withholding treatment (WW) and control (C) in three replicates. Single row (7 plants) was planted for each genotype in each treatment in each replicate. Watering regime was optimized in preliminary trials to obtain the 50% reduction in fresh weight per plant in WW treatment compared to C. Plants in C were watered with spray bottle in planting and every two days thereafter with 8 ml of tap water per plant. Plants in WW were watered in planting with full dose (8 ml) and two times thereafter (last watering on 4^th^ day) with half a dose of water added per plant in C (4 ml). After that, water was withheld to 14^th^ day when the aboveground parts of three equally developed plants per genotype in each replicate were harvested and prepared for further analysis. Plants were weighed on a precise four decimal scale to obtain fresh weight (FW) and the 1 g samples were chopped, put in previously weighed 10 ml Falcon tubes and weighed together. Tubes with samples were left for 24 h in digital laboratory drying oven on 80°C, and weighed. Dry weight (DW) was calculated from product of FW and dry weight (sample)/fresh weight (sample). Dry matter content (DMC) was calculated as (DW/FW) × 100. Additionally, the hundred kernel weight (HKW) was estimated by weighing five random samples of hundred kernels mixed from five self-pollinated ears.

### 2.2. Phenotypic data analysis

Phenotypic data were analyzed with mixed linear model in *lme4* library [21] using R programming language [22]. Repeatabilities within the treatments were estimated from variance components as *R^2^* = σ^2^_G_ / (σ^2^_G_ + σ^2^_e_/r), while the heritability of each trait was estimated as *H^2^* = σ^2^_G_ / (σ^2^_G_ + σ^2^_GxT_/t + σ^2^_e_/tr) where G represents genotypic effects, GxT is genotype-treatment interaction, e is error and t and r are the numbers of treatments and replicates respectively. *Post-hoc* tests were performed using the R/*agricolae* package [23] and generalized linear model.

### 2.3. GBS data and filtering

The GBS genotyping was performed on Cornell University [14] according to protocol developed by Elshire et al. [24] and the data were obtained from PANZEA organization [25] repository. The original dataset contained 945,574 SNPs that were annotated, partially imputed, and assigned to chromosomes. Filtering of the low quality SNPs was performed with Tassel software [26] version 5.2.5. Namely, maximum of heterozygotes was set to 0%, minor allele frequency to 2.5%, and maximum of missing data per position to 5%. After filtering there were 229,680 SNPs left. To avoid redundancy of SNPs in similar positions, thinning to 1 SNP per 1,000bp was carried out leaving totally 40,777 high quality markers.

### 2.4. Genetic structure analysis and linkage disequilibrium

The genetic structure present in the collection was determined with software STRUCTURE, version 2.3.4 [27]. Random sample of 10,000 SNPs was taken as the input for genetic structure analysis. Analysis was carried out with 10,000 burn-in cycles and 50,000 MCMC runs and population admixture assumed. STRUCTURE analysis was set with 10 populations (K = 10) and four runs were carried out for each value of K. The best number of K was 2 and it was chosen according to ΔK method [28] using CLUMPAK software [29]. The germplasm set was divided into groups corresponding to Stiff Stalk (SS) and Non-Stiff Stalk (NS) derived germplasm. Additional run with 50,000 burn-in cycles and 100,000 runs was carried out to generate the Q matrix. The Q matrix is available in S1 Table.

The linkage disequilibrium (LD) decay with physical distance was calculated using Tassel software with window size of 50 bp comprising totally 2,622,025 calculations. The LD decay plot has been constructed in R using the local regression smoothing function *loess*. Across all chromosomes, LD decayed at 0.12 Mbp.

### 2.5. Marker effects and GWAS

Mixed linear modelling approach with structure (Q) and kinship (K) matrices (MLM+Q+K) was used for GWAS analysis to control for spurious associations and false positives. Analyses were performed with Tassel software version 5.2.5. The Q matrix for MLM was calculated using all 40,777 SNP markers with principal component analysis with two axes according to the genetic structure results. Correlation between Q values assessed by STRUCTURE method and eigenvectors from PC analysis was 0.98. The phenotypic data provided for association mapping were plot values with replicates set as covariates. Two −log(*P*) value thresholds for controlling the false detection rate (FDR) were applied. The value of the first threshold was determined according to the Bonferroni correction for α=0.05 significance level. Namely, the alpha value (α=0.05) was divided by the effective number (M_*eff*_) of 14,155 SNP markers and the value of 5.4519 was used as threshold for declaring the significantly associated loci. The value of M_*eff*_ was calculated according to multiple testing method as implemented in *SimpleM* R script [30, 31]. According to the results of Bian and Holland [32] that showed the stable predictive abilities of the loci detected in the range of −log(*P*) thresholds from 4 to Bonferroni corrected value in oligogenic and polygenic traits, another less-stringent threshold of 4 was applied.

The correlations between the marker effects were calculated according to Ziyomo and Bernardo [33] and Galic et al. [34] to estimate the similarity of small-effect loci governing the different traits even if they fall below the thresholds set for GWAS. The marker data for calculation of ridge regression best linear unbiased predictions of marker effects (rrBLUP) were re-coded with Plink software version 1.07 [35] and R interface. The rrBLUP model was set with *rrBLUP* R library [36] using the library mixed-model solver integrated in *mixed.solve* function with markers as random effects and fixed intercept. Correlations between BLUPS of marker effects among different traits were calculated.

Candidate genes were counted through the interface of Maize GDB webpage [37]. The analysis of candidate genes was limited to protein-coding genes within the same linkage disequilibrium block (*R^2^*<0.2) of 120 kbp around the significantly associated SNP. Physical locations of SNPs are reported according to Maize B73 RefGen_v4 map.

## 3. Results

### 3.1. Genetic structure and linkage disequilibrium

According to the ΔK method, two genetic subgroups were detected in the STRUCTURE analysis (Figure 1a). In alignment with the Q values of the reference lines B73 and Mo17, the two subgroups were designated SS and NS for brevity. PC analysis indicated higher diversity in NS compared to SS subgroup (Figure 1b). The Q matrix for all 109 lines used in this study is available in S1 Table. Linkage disequilibrium decayed across all 10 chromosomes below the arbitrary threshold of *R^2^*<0.2 at 120 kbp distance between pairs of markers (Figure 1c).

**Figure 1.**
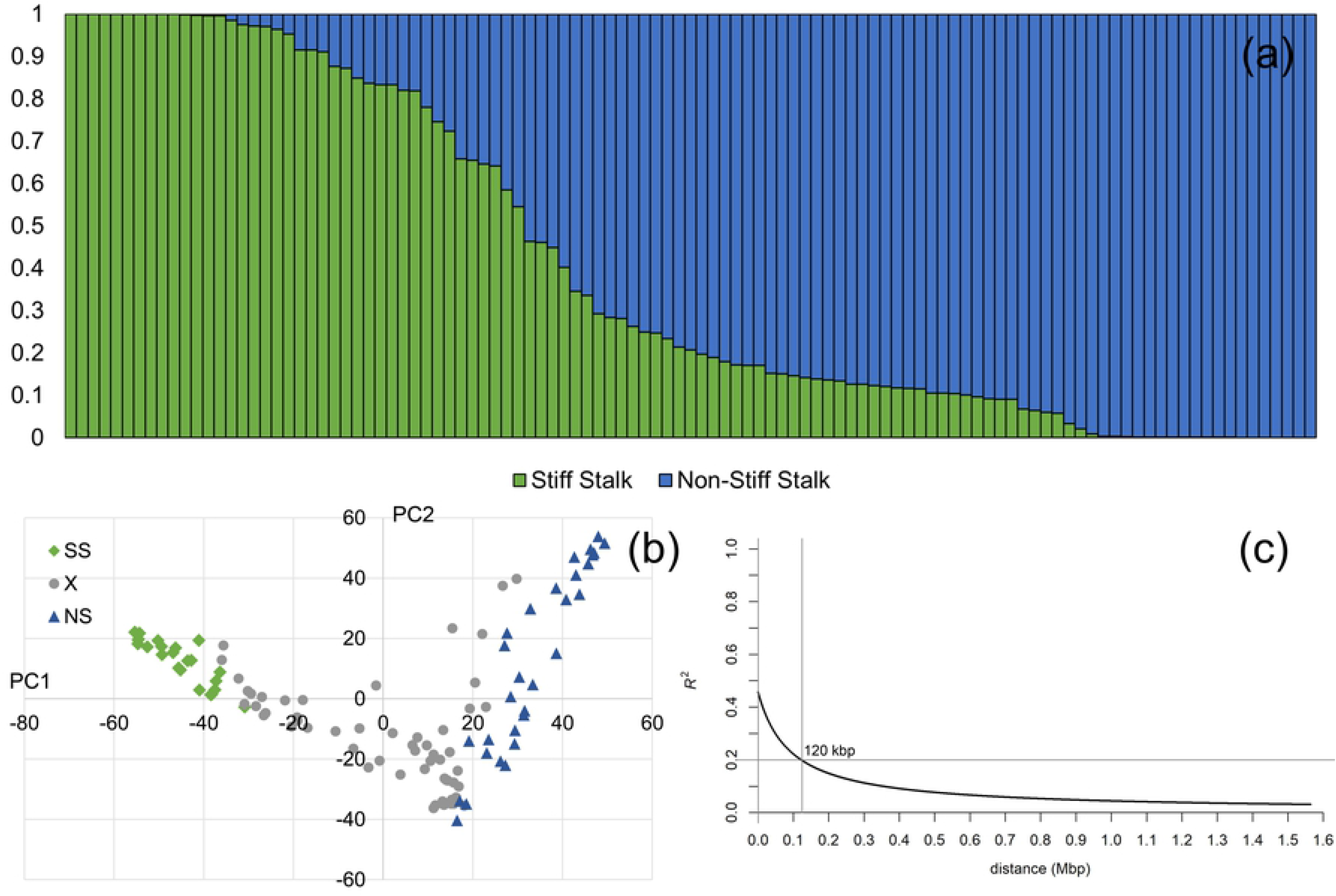
Genetic properties of the 109 maize inbred lines with expired PVP certificate: (a) Q matrix from STRUCTURE analysis; (b) results of PC analysis using 40,777 GBS SNPs – marked green and blue are the lines with Q>0.9 for Stiff Stalk (SS) and Non-Stiff Stalk (NS) groups, respectively (lines in gray denoted by X are of mixed origin); (c) linkage disequilibrium decay across all chromosomes

### 3.2. Variance components, heritabilities and responses to water withholding

The significant effects of WW treatment were detected for all three examined traits (Table 1). Effects of water withdrawal manifested as significant 41.3% reduction in mean FW, 19.5% reduction in mean DW and 39.7% increase in DMC relative to C. High estimates of genetic variance were observed for all examined traits (Table 1). Interestingly, while the genetic variances of FW and DW have decreased in WW treatment, the genetic variance of DMC increased 4.4 times in WW. Larger estimate of genotype-treatment interaction (72% of σ^2^_G_) was detected for DMC compared to FW and DW in combined analysis. Very high repeatabilities and heritabilities were determined for all examined traits. The cause of high repeatability values of HKW, FW and DMC were low obtained estimates of error variances in addition to the high estimated values of σ^2^_G_. Heritability estimates of FW, DW and DMC across both treatments were expectedly lower due to the accounting for genotype-treatment variance.

**Table 1.** Means ± standard deviations, ranges, variance components, repeatabilities and heritabilities for hundred kernel weight (HKW) in grams and biomass traits fresh weight (FW) in grams, dry weight (DW) in milligrams and dry matter content (DMC) as % of FW in control (C) and water withholding (WW) treatments. The values calculated within treatments are repeatabilities, while the values calculated across treatments represent heritabilities. Repeatability of HKW is calculated for year of seed multiplication (2018).

| Trait | Treatment | Mean ± SD<sup>a</sup> | Range | $\sigma^2_G$ | $\sigma^2_{G \times T}$ | $\sigma^2_e$ | $H^2$<sup>b</sup> |
| --- | --- | --- | --- | --- | --- | --- | --- |
| HKW (g) | – | 24.91 ± 4.07 | 14.68 – 38.40 | 14.79 | – | 1.90 | 0.96 |
| FW (g) | C | 0.92 ± 0.26 a | 0.31 – 1.49 | 0.05 | – | 0.02 | 0.88 |
| | WW | 0.54 ± 0.18 b | 0.11 – 1.00 | 0.02 | – | 0.01 | 0.85 |
| DW (mg) | C | 60.29 ± 19.71 a | 13.78 – 106.07 | 265.00 | – | 124.10 | 0.86 |
| | WW | 48.57 ± 17.45 b | 11.10 – 87.62 | 199.00 | – | 107.40 | 0.85 |
| DMC (%) | C | 6.51 ± 1.05 b | 3.99 – 10.26 | 0.55 | – | 0.50 | 0.77 |
| | WW | 9.10 ± 1.86 a | 3.96 – 13.16 | 2.43 | – | 1.05 | 0.87 |
| FW (g) | Combined | 0.73 ± 0.29 | 0.11 – 1.49 | 0.03 | 0.01 | 0.02 | 0.80 |
| DW (mg) | Combined | 54.44 ± 19.51 | 11.10 – 106.07 | 210.72 | 21.11 | 116.16 | 0.88 |
| DMC (%) | Combined | 7.80 ± 1.98 | 3.96 – 13.16 | 0.86 | 0.63 | 0.78 | 0.66 |

<sup>a</sup> different letters denote significant differences between treatments according to LSD test at $\alpha=0.05$

Responses of the two genetic subgroups (SS and NS) of inbreds were further analyzed to identify the sources of favorable alleles for responses to water withholding in this stage (Figure 2). Inbreds were split to SS and NS groups according to their respective results from STRUCTURE analysis. Only responses of lines with ancestry coefficient >0.9 were analyzed in this regard. More diverse responses to WW were observed in NS group compared to SS group which was probably caused by inability of STRUCTURE method to infer ancestry of other genetic groups due to insufficient number of samples with unmixed pedigrees in NS pool. Interestingly, lines PHW65 and LH38 from NS pool with related pedigrees showed stable FW values in WW, accompanied with increase in DW. Increase in DW accompanied by decrease in FW was also observed in related lines LH132, LH195 and LH205 from SS pool. The largest relative increase in DMC was observed in line C103 (3.99% in C, 9.84% in WW).

**Figure 2.**
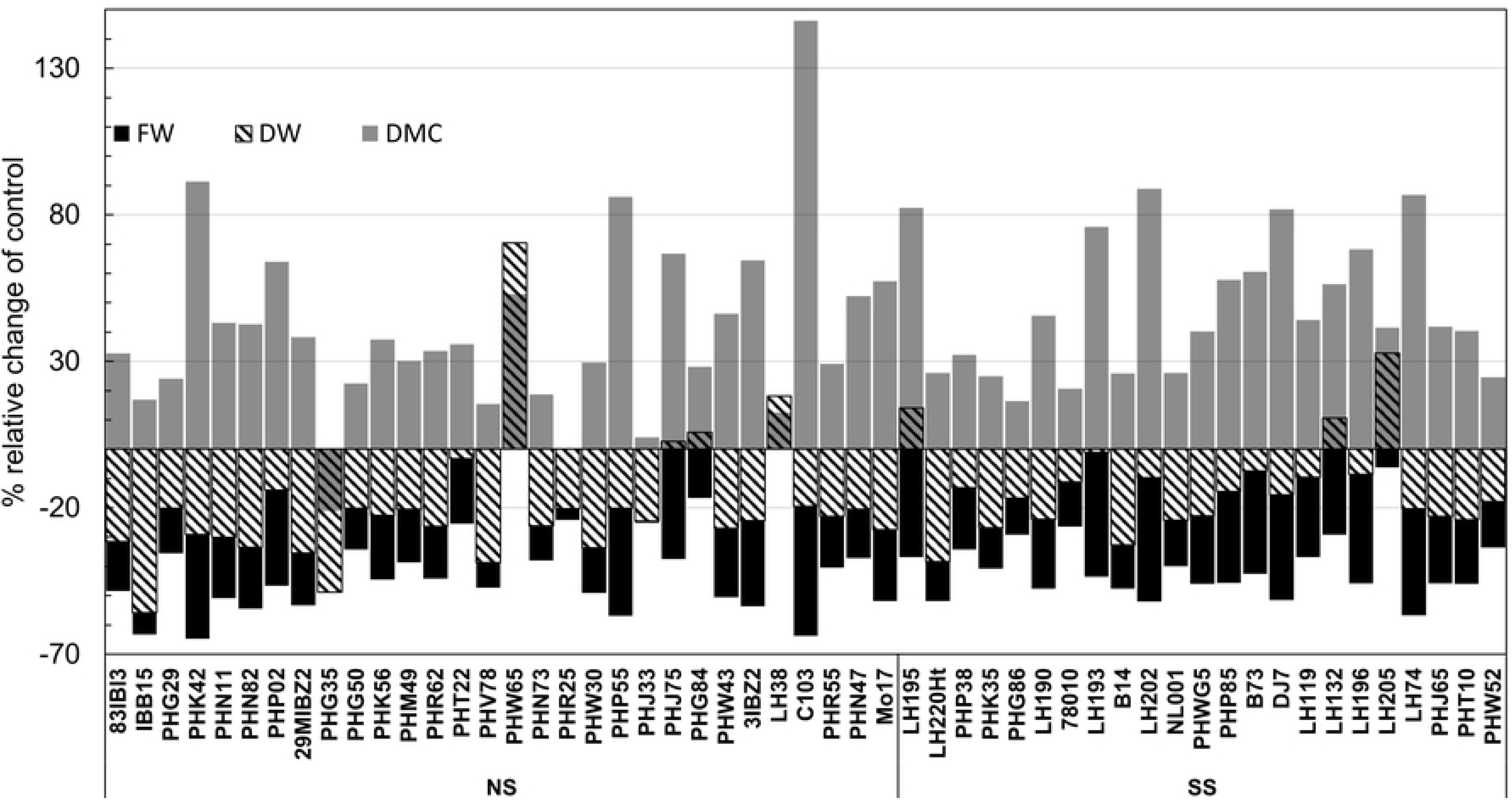
Relative changes in fresh weight (FW), dry weight (DW) and dry matter content (DMC) in water withholding (WW) treatment compared to control (C) of inbred lines with Q>0.9 to SS or NS genetic group (Figure 1a).

Strong phenotypic correlations were observed for FW and DW between treatments, while correlation of DMC between the treatments was moderate (Figure 3, upper triangle). Correlations between FW and DW were strong to very strong within and between C and WW. Weak to moderate correlation was observed between DW and DMC across the treatments, while there were no significant correlations detected for FW and HKW with DMC. Weak to moderate correlations were detected for FW and DW with HKW in both C and WW. Correlations of rrBLUP marker effects (Figure 3, lower triangle) clearly resembled phenotypic correlations, which reflects the high estimates of genetic variances for the traits within the treatments and low error estimates (Table 1). Near zero estimate of correlation of marker effects between DMC and FW indicated that there is no expected shared genetic background between the traits. Correlation of DMC across the treatments was moderate. Weak correlations between FW, and DW in WW with DMC indicated different mechanisms in genetic regulation of these traits.

**Figure 3.**
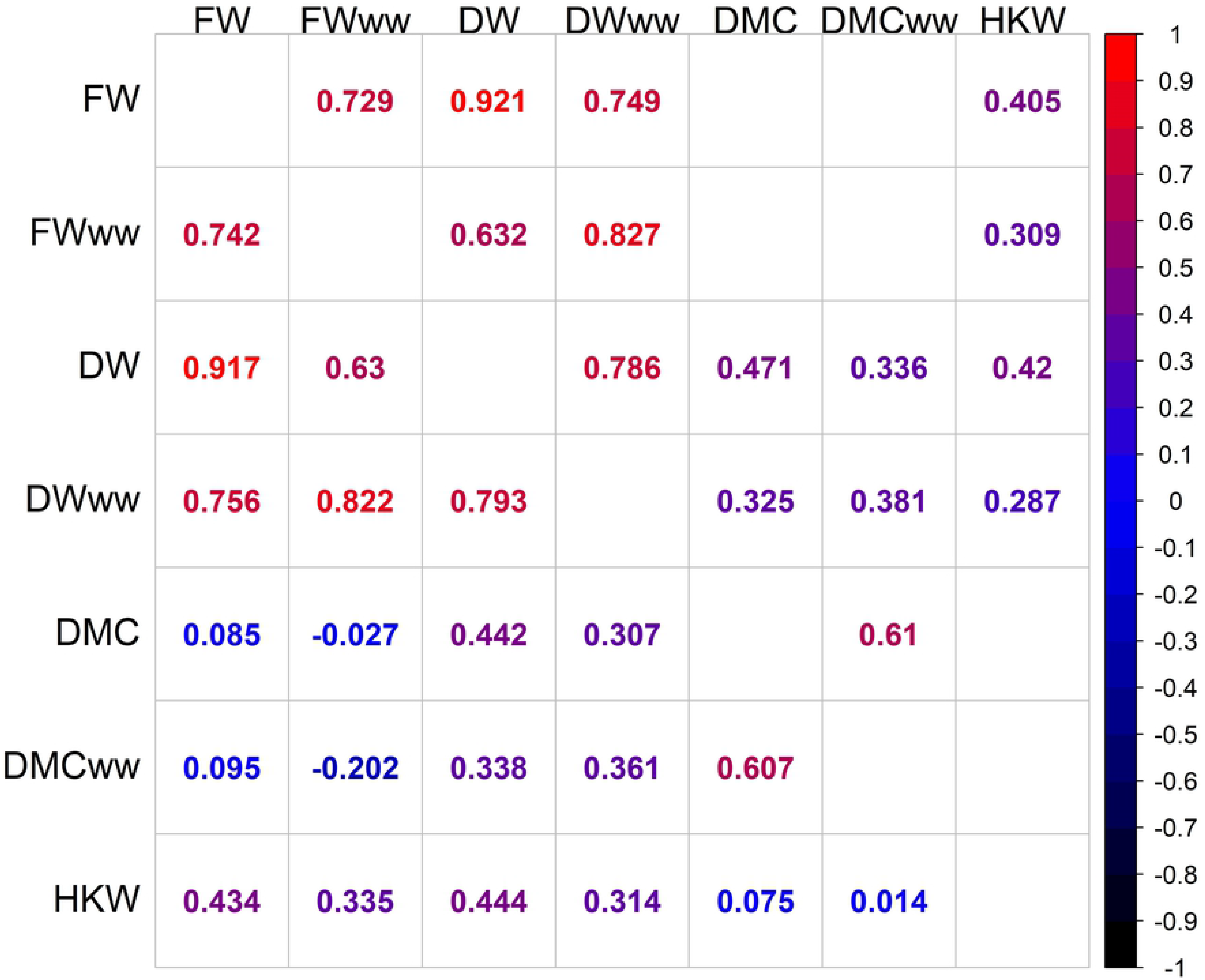
Pearson’s correlation coefficients (upper triangle) among fresh weight (FW), dry weight (DW), dry matter content (DMC) between control (C) and water withholding (WW) treatments and with hundred kernel weight (HKW). Lower triangle represents Pearson’s correlations between rrBLUP estimates of marker effects between traits. Shown are the correlations significantly different from zero at α=0.05.

### 3.3. Allelic effects and candidate genes

Inspection of QQ plots showed there were no highly significant SNP effects detected for FW in C and WW, DW in C and DMC in WW in MLM+Q+K association mapping procedure (Figure 4). Positive deviations from the expected distribution of −log(P) values were observed above value of 3 for DMC in C and above 4 for DW in WW. The total number of loci crossing the arbitrary – log(P) threshold of 4 was 29 (Figure 5), while only 3 loci for DMC in C crossed the calculated Bonferonni threshold of 5.45 (Figure 5e). No SNPs crossing the arbitrary threshold of 4 was detected for FW in C. The detected loci were distributed over 6 of 10 chromosomes (1, 2, 3, 6, 7, 9). *R^2^* values of the detected loci ranged from 4.71 to 9.33% (Table 2). Single pleiotropic locus was detected for FW and DW in WW on chromosome 2, position 17.341 Mbp. A QTL on chromosome 6, position 101.971 Mbp detected for DMC in C was also detected in WW. QTL for DMC in C on chromosome 1, position 175.378 is probably the same as the QTL for DMC in WW on position 170.174, or it might represent another tightly linked QTL.

**Figure 4.**
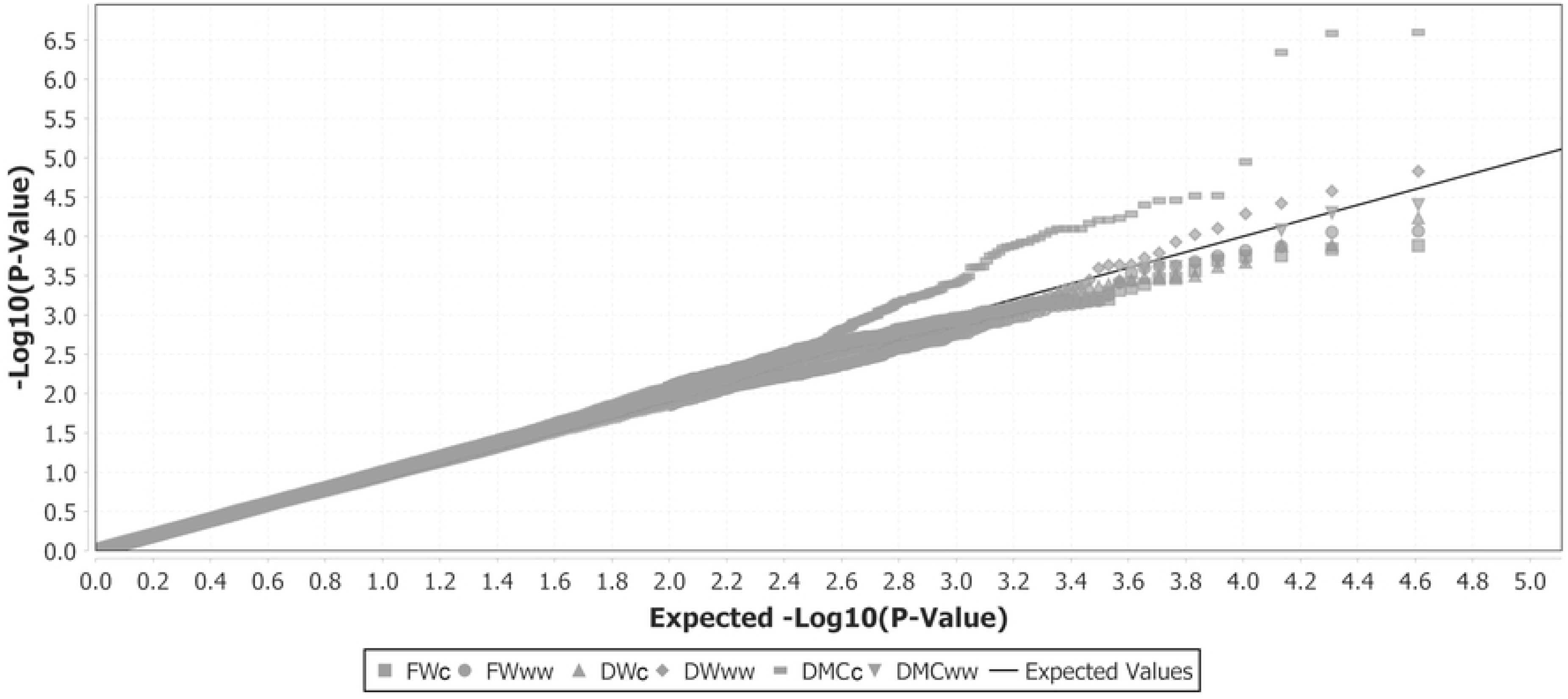
The quantile-quantile plot of the −log(P) values of the allelic effects from MLM+Q+K analysis for fresh weight (FW), dry weight (DW), dry matter content (DMC) between in control (c) and water withholding (ww) treatments.

**Figure 5.**
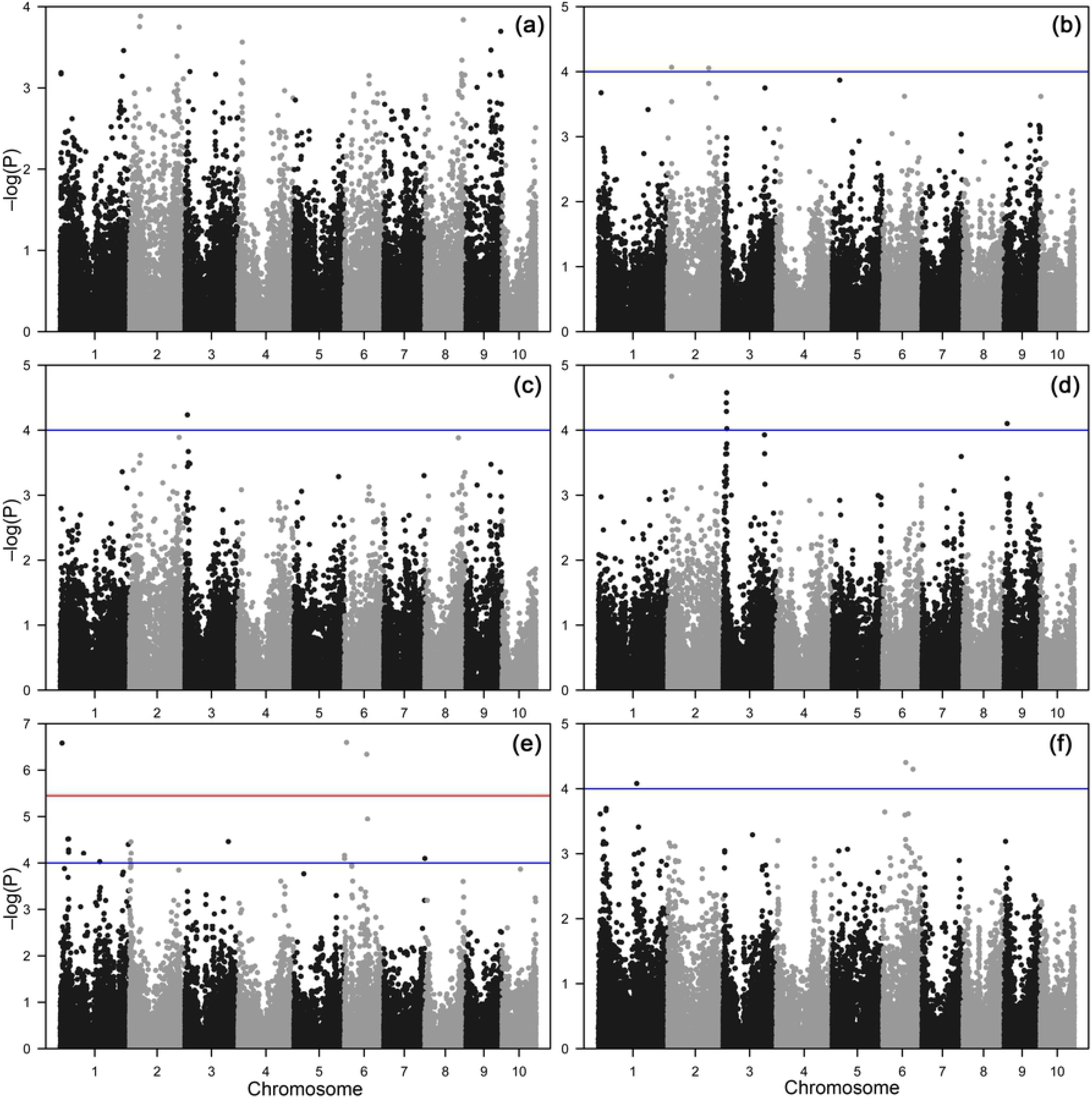
The Manhattan plots of the −log(P) values of the allelic effects from MLM+Q+K analysis for fresh weight in control (a), fresh weight in water withholding treatment (b), dry weight in control (c), dry weight in water withholding treatment (c), dry matter content in control (e) and dry matter content in water withholding treatment (d). Blue line represents arbitrary threshold value of 4, while the red line represents the Bonferroni corrected threshold value of 5.45.

**Table 2.**
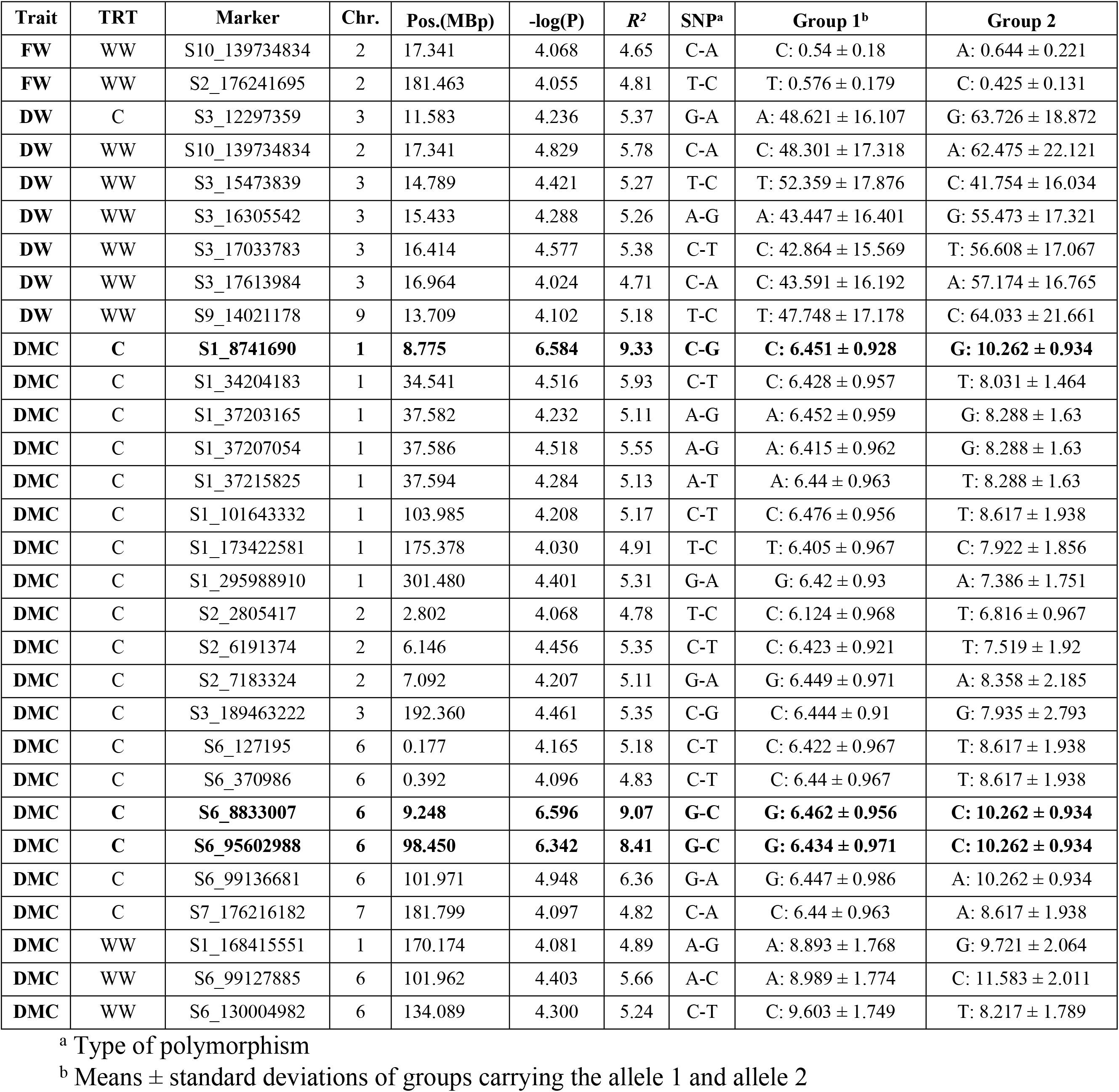
SNPs crossing arbitrary threshold of 4 associated with biomass traits fresh weight (FW) in grams, dry weight (DW) in milligrams and dry matter content (DMC) as % of FW in control (C) and water withholding (WW) treatments. SNPs in bold are the ones crossing Bonferroni corrected threshold value for significance at α=0.05.4.

## Discussion

### 4.1. Genetic structure

The results of PC analysis in this study were similar to those of Beckett et al. [38] in a study of genetic diversity in ex-PVP germplasm. In their study, STRUCTURE method combined with ΔK population number selection procedure was able to differentiate three independent clusters, namely SS, NS and Iodent, while in our study Iodent, although visually present, was confounded to NS pool due to insufficient number of inbreds with unmixed origin from two putative NS subgroups. Linkage disequilibrium was higher compared to large diversity panels [14,38] but comparable to recent reports with medium sized maize diversity panels [39,40].

### 4.2. Responses to water withholding

Fresh and dry matter accumulation in early vegetative stages of growth is heavily influenced by water deficit, and phenotyping for these traits in diverse germplasm panels under water deficit is one of the first steps in understanding the drought tolerance [41]. Avramova et al. [20] showed that biomass traits such as FW, DW and DMC are valuable in monitoring responses to water deficit, and can feasibly differentiate even the slight differences in responses of different cultivars. The drought treatment in their study was more severe (14 day WW) which also changed the scale of responses in all traits and the plants were analyzed in different stage compared to this present study (V4 compared to V3). However, the directions of responses noted were the same. This is also in accordance with the results of Sekhon et al. [42] that showed how genetic mechanisms for stress responses regulation appear to be related to their respective tissues, and so the leaf transcriptome changes in limited manner from the first leaf appearance to the end of vegetative growth phases. Ge et al. [43] also used these traits for phenotyping two maize inbred lines differing in tolerance to water deficit. It was shown that different responses are expected very shortly after the water withholding, while the effects of WW after 9 days of treatment started to become more pronounced. Authors also found that these traits can be efficiently predicted using the hyperspectral reflectance indices. Most interestingly, two inbreds from NS pool in our study considerably increased their DW in WW treatment. Such responses to WW by increase in DW were reported in literature for drought resistant maize inbred lines [44,45]. The accumulation of DW in WW was probably induced by accumulation of osmolytes, and increased activities of antioxidant pathways. The higher diversity of reactions in NS pool compared to SS pool probably reflected the genetic diversity confounded within the NS pool (S1 Table).

The choice of mixed modelling approach in variance components analysis was supported by results of van Eeuwijk et al. [46] that more and more researchers use mixed models to precisely address the GxE variance. The GxT variance in our study is also a form of GxE variance, although with strictly specified conditions of E and changes compared to reference environment (C). The very high repeatabilities of traits examined in this study were expected for experiments set in the controlled conditions addressing the quantitative traits.

The low estimates of GxT interaction for FW and DW were accompanied by high levels of plasticity in reactions of examined genotypes for these traits. Plasticity was detected through the correlation analysis in which performances of FW and DW in C were strongly correlated to performances in WW. In contrast, considerable proportion of GxT variance for DMC accompanied by lower correlation estimates for performance between treatments indicated the presence of different mechanisms of coping with water deficit, and divergence in responses of different genotypes to stress [47]. Traits FW and DW were moderately correlated to HKW in control, and weakly in WW. This was supported with the findings that early vigor of maize plants is influenced by kernel weight [48], but variations in kernel size appear to offer no clear advantage when water is sparse [49].

### 4.3. Allelic effects and candidate genes

In correlation analysis of marker effects (Figure 3, lower triangle) according to the studies of Ziyomo and Bernardo [33] and Galic et al. [50], it was shown that marker effects for FW in C explain ~56% of variance for FW in WW. The marker effects for DW in C explain ~63% variance for DW in WW, while the marker effects for DMC in C explain only 37% variance of marker effects for DMC in WW. Interestingly, marker effects for FW in WW treatment explain ~68% variance of marker effects for DW in WW treatment. It is expected that FW and DW are highly correlated, as accumulation of osmotically active compounds into the aboveground plant parts is a strategy contributing to increase in both FW and DW in water-limited conditions [20]. The high estimates of correlations between the rrBLUP marker effects for traits between treatments show there is much of the overlap between the genetic mechanisms regulating these responses, although small number of loci affecting same traits between treatments was detected in GWAS. This is partially explained by the findings that the allelic effects for quantitative traits are highly sensitive to environmental changes that could be attributed to different scenarios [51]. Certain alleles affecting quantitative traits can decrease in effect size in conditions when water is in deficit [52] which could be the reason for the small number of significant allelic effects between treatments despite the high correlations of rrBLUP marker effects. In addition to that, the limiting factor in detection of loci is still relatively strict threshold of 4 applied in association analysis compared to arbitrary thresholds found in literature.

The GWAS analyses of responses to relevant stresses using relevant phenotypes such as plant’s biomass traits provide information not only valuable in that particular regard, but also in wider perspective of plant’s stress responses, as the plant’s genetic mechanisms of coping with different stresses are similar to certain extent [53]. The search for candidate genes was limited to loci which crossed the Bonferroni threshold, the loci with significant effect across the treatments and putative pleiotropic loci on chromosomes 1, 2 and 6. Considering genes located within the estimated 240 kbp window (± estimated linkage disequilibrium block of 120 kbp) from loci crossing Bonferroni threshold (all for DMC in C), 11 candidate genes were counted for the QTL on chromosome 1, position 8.775 Mbp, 1 candidate gene for the QTL on chromosome 6, position 9.248 Mbp, and 7 for the QTL on chromosome 6, position 98.450 Mbp. A putative cytochrome P450 superfamily protein is identified for DMC in C on chromosome 1. Cytochrome P450 monooxygenases mediate the biosynthesis of various secondary compounds that act as plant defense agents [54]. Metabolic studies and mRNA analyses have suggested individual P450s or subsets of these P450s are induced in response to chemical inducers and varying environmental conditions [55]. On chromosome 6, position 9.248 Mbp, single candidate gene was found. The gene LSD1 coding for protein lesion simulating disease 1 was shown to interact with cellular antioxidant defense mechanisms, namely catalase, thus causing the ROS build-up caused by functions of peroxisomes [56]. The LSD1 – catalase interaction also has an important role in control of the programmed cell death. N-acetyltransferase HLS1, identified as a putative candidate gene for DMC in C on chromosome 6, position 98.450 Mbp is known to control differential cell growth by regulating auxin activity in *Arabidopsis* [57]. Also, auxin signaling pathways were shown to be differentially regulated in inbred lines differing in drought susceptibility [58]. Three candidate genes were identified for putative pleiotropic locus detected affecting FW and DW in WW on chromosome 2. One of them is the calmodulin binding protein known to be included in stress responses of plants [59]. The role of calmodulin in plants is the maintenance of homeostasis between different cellular processes and prevention of interference between components of different biochemical mechanisms which is especially pronounced during the plant responses to stress. In the region 70 kbp upstream from the QTL for DMC in both WW and C on chromosome 6, position 101.97 Mbp, gene coding for enzyme polygalacturonase is found. Polygalacturonases are endogeneous pectinases usually expressed in expanding tissues and guard cells and have important function in young plant development [60], which can be linked to the trait DMC in our study, that provides the information regarding the dry matter accumulation regardless of the treatment.

## Conclusions

In this study it was shown that biomass traits which are easily measured and show high heritabilities can be used to assess the responses of young maize plants to water withholding. The similar genetic mechanisms were found controlling traits FW and DW in both C and WW. Similarity of the genetic mechanisms was more visible through the strong correlations of rrBLUP marker effects between the traits than through the results of GWAS, which was probably caused by relatively stringent threshold of 4 compared to the thresholds reported in the literature and the variation in allelic effects of some loci affected by the water deficit [52]. Interestingly, the biparametric expression DMC calculated from FW and DW was shown to have properties of a completely independent phenotype. Furthermore, we detected the highest GxT interaction for DMC and the differentiation in genetic mechanisms regulating DMC between the treatments. The use of biomass traits in breeding tolerance to water withholding in early vegetative stages of growth appears to offer cost-effective, data-rich approach, but more studies are needed linking these traits to other physiological processes thus facilitating the build-up of understanding the processes involved in abiotic stress responses.

## Acknowledgements

This work has been supported by the EU project KK.01.1.1.01.0005 “Biodiversity and Molecular Plant Breeding” of the Centre of Excellence for Biodiversity and Molecular Plant Breeding (CroP-BioDiv), Zagreb, Croatia.

## Supporting information

**S1 Table.** Publicly available information for the inbred lines used in the experiments, along with Q values from the STRUCTURE analysis with 50000 burn-in cycles and 100000 MCMC replicates

**S2 Figure.** The cumulative precipitation in Osijek, Croatia mapped to theoretical percentiles of the 50-year historical data for 2015 (a), 2016 (b), 2017 (c) and 2018 (d). Source: Croatian Hydrological and Meteorological service, Republic of Croatia

